# A Study of Event-Related Potentials during Monaural and Bilateral Hearing in Single Sided Deaf Cochlear Implant Users

**DOI:** 10.1101/2022.06.14.495873

**Authors:** Marcus Voola, An T. Nguyen, Andre Wedekind, Welber Marinovic, Gunesh Rajan, Dayse Tavora-Vieira

## Abstract

**Objective:** Single sided deafness (SSD) is characterized by a profoundly deaf ear and normal hearing in the contralateral ear. A cochlear implant (CI) is the only method to restore functional hearing in a profoundly deaf ear. In a previous study, we identified that the cortical processing of a CI signal differs from the normal hearing ear (NHE) when directly compared using an auditory oddball paradigms consisting of pure tones. However, it is unclear how the electrical and acoustic signals from each ear are combined. This study aims to understand how the provision of the CI in combination with the NHE may improve SSD CI users’ ability to discriminate and evaluate auditory stimuli.

**Design:** Electroencephalography (EEG) from ten SSD-CI participants (four participated in the previous pure-tone study) were recorded during a semantic acoustic oddball task, where they were required to discriminate between odd and even numbers. Stimuli were presented in four hearing conditions: directly through the CI, directly to the NHE, or in free field with the CI switched on and off. We examined task-performance (response time and accuracy) and measured N1, P2, N2N4 and P3b event-related brain potentials (ERPs) linked to the detection, discrimination, and evaluation of task relevant stimuli. Sound localization and speech in noise comprehension was also examined.

**Results:** In direct presentation, task performance was superior during NHE compared to CI (Shorter and less varied RTs [∼720 vs. ∼842 ms], higher target accuracy [∼93 vs. ∼70%]) and early neural responses (N1 and P2) were enhanced for NHE suggesting greater signal saliency. However, the size of N2N4 and P3b target-standard effects did not differ significantly between NHE and CI. In free field, target accuracy was similarly high for both FF-on and -off (∼95%), with some evidence of CI interference during FF-on (more variable and slightly but significantly delayed RTs [∼737 vs. ∼709 ms]). Early neural responses and late effects were also greater during FF-on. Performance on sound localization and speech in noise comprehension (S_CI_N_NHE_ configuration only_)_ was significantly greater during CI-on.

**Conclusion:** Both behavioral and neural responses in the semantic oddball task were sensitive to CI in both direct and free-field presentations. Direct conditions revealed that participants could perform the task with the CI alone, although performance was sub-optimal and early neural responses was reduced when compared to the NHE. For free-field, the addition of the CI was associated with enhanced early and later neural responses but did not improved task performance. Enhanced neural responses show that the additional input from the CI is modulating relevant perceptual and cognitive processes, but the benefit of binaural hearing on behavior may not be realized in simple oddball tasks which can be adequately performed with the NHE. Future studies interested in binaural hearing should examine performance under noisy conditions and/or employ spatial cues to allow headroom for the measurement of binaural benefit.

## Introduction

### Single Sided Deafness

Single sided deafness (SSD) is a condition characterised by a severe to profound sensorineural hearing loss in one ear (> 70 dB HL) and normal hearing in the other (< 20 dB HL)(Sullivan et al. 2020). The condition is estimated to affect ∼ 0.1 – 0.5 % of the population (Golub et al. 2018). The reliance on only one functioning ear results in reduced localisation accuracy and increased difficulty understanding speech in noise (Galvin et al. 2019). A cochlear implant (CI) can re-establish this loss of input by directly stimulating the auditory nerve in the impaired ear, potentially regaining binaural hearing. Improvements in binaural hearing after CI has been noted in past studies which showed that the provision of a CI improves speech in noise comprehension and localisation ability (Peter et al. 2019; Purdy et al. 2016; Tavora-Vieira et al. 2015; Tokita et al. 2014; Vermeire et al. 2009; Wedekind et al. 2020).

Despite the CI’s ability to restore hearing in the impaired ear, the electrical signal produced by the CI does not contain all the spectral information that the normal hearing ear (NHE) provides. Consequently, it provides the user with a reduced amount of auditory information compared to the information received by the NHE (Drennan et al. 2008). Additionally, for the benefits of binaural hearing to be reaped, the brain needs to integrate the acoustic signal from the NHE with the electrical signal from the CI. Currently, it is not completely understood how the brain uses the electrical signal from the CI – independently or in conjunction with the NHE. One method of examining these neural processes is by using electroencephalography (EEG) and the event-related potential (ERP) technique to measure how cortical brain areas respond to auditory stimuli across listening scenarios (e.g., CI vs. NHE).

### Examining Auditory Processing using Event-Related Potentials

EEG is used to study changes in cortical activity immediately following an event of interest (e.g., presentation of a sound/image, or onset of a button-press) via an array of scalp electrodes. The ERP technique is an analysis method that quantifies the amplitude, latency and spatial distribution of peaks induced by an event, which offers an insight into the different stages of auditory processing – such as the initial detection and subsequent high-order processes associated with interpreting the stimulus (Light et al. 2010). ERPs can be elicited by an oddball paradigm that consists of a frequent (standard) and infrequent (target) stimulus, whereby the target stimuli are characterised by a unique feature (Polich 2007). Initial studies typically used pure tones with differing frequencies (e.g., 1 vs. 2 kHz) or different speech tokens (e.g., /ba/ vs. /ka/) and focused on measuring four ERP components: the N1, P2, N2N4 and P3b (Didoné et al. 2016).

The N1 and P2 refers to frontocentral negative deflection occurring ∼ 100 ms and a frontocentral positive deflection occurring ∼150 ms after stimulus onset. Both deflections have been implicated in stimulus-detection and attention orientation based on its early timing (Legris et al. 2018; Tremblay et al. 2014). Modulation in amplitude and latency are interpreted as reflecting efficiencies in stimulus-detection and/or attention orientation.

In auditory oddball paradigms, research is generally focused on later ERP components such as the N2N4 and P3b. The N2 refers to a frontocentral distributed negativity occurring ∼200-350 ms after stimulus onset and is thought to reflect stimulus-discrimination as it is typically enhanced in amplitude for novel or task-relevant stimuli compared to frequent or irrelevant stimuli (Lau et al. 2008; Näätänen et al. 2011). In tasks with complex stimuli (e.g., words), this N2 is typically delayed and exhibits a broader time-course (∼400-600 ms) and is sometimes referred to as the N2N4 – highlighting its variation from the N2 (Song et al. 2018).

The P3b refers to a parietally distributed positivity which occurs around 300–600 ms after stimulus onset (Polich, 2007). Like the N2, this component is enhanced for novel and task-relevant stimuli compared to irrelevant stimuli. Based on its later time-course, the P3b is thought to reflect deeper evaluation of stimuli such as context-updating, which is the updating of stimulus probabilities in memory (Donchin 1981; Polich 2007). More recently, the P3b is thought to represent the decision making process, reflecting stimulus categorisation and the activation of stimulus-response links (Verleger et al. 2014).

### ERPs in Single Sided Deaf Cochlear Implant users

Several studies have examined brain potentials of SSD-CI users during a variety of tasks, under different configurations. These include (1) the passive listening of speech-tokens in free-field, in relation to behavioural metrics such as sound localisation and speech-in-noise, (2) active discrimination of pure tones in an oddball task via direct connect and headphones (Wedekind et al. 2021), and (3) active discrimination of words during free field (Finke et al. 2016).

Recently, one ERP study has examined P2 in SSD CI users during a passive listening of speech-tokens (/ba/) in free-field listening. In a between-groups study, Legris and colleagues (2018) reported reduced P2 amplitudes in experienced SSD CI users compared to normal hearing controls, concluding that the ability to detect sounds during bilateral listening was only partially restored (Legris et al. 2018). One difficulty with comparing raw voltages in between-subject data is that inter-individual differences in skull-thickness, the location and orientation of dipoles, and variation in impedances values may influence neural responses. While partial restoration of bilateral hearing is consistent with functional assessments (Galvin et al. 2019; Távora-Vieira et al. 2015), it is difficult to assess contribution of CI from other factors.

Another approach is to compare neural responses between ears (i.e., CI vs. NHE) within the same subject. When we compared auditory detection between CI and NHE, we observed no difference in P2 latency, suggesting that the detecting processes are operating similarly (Wedekind et al. 2020). While neural responses are similar between CI and NHE, the provision of the CI leads to clear improvements in spatial acuity and speech-in-noise performance (Snapp et al. 2017; Távora-Vieira et al. 2015; Williges et al. 2019), highlighting a gap in our understanding of the brain-behaviour relationship. One way to assess the functional significance of these components, is to use an active discrimination task and examine how brain responses during different listening conditions are associated with changes in reaction time and accuracy.

In a recent study, our lab has directly contrasted CI and NHE during active discrimination of pure tones using an oddball task. We found that the CI led to benefits in sound localisation and speech-in-noise – in line with previous work. Individually, we identified that the brain signals from NHE and CI were capable of discriminating target and standard tones, but there were slight differences in efficiencies. (Wedekind et al. 2021). In 2017, Finke et al (2017) further investigated the higher order processing capabilities within the SSD CI population by employing the use of a semantic oddball paradigm. SSD CI participants were required to classify words as being either living or non-living entities, therefore differentiating stimuli on more than just the physical properties. Using more complex stimuli is beneficial to this field as it may overcome any ceiling effects that are present with simple oddball task and thereby revealing any deficits in the processing of auditory stimuli. It was identified that the latency of the N2N4 and P3b was delayed for the CI ear when compared with the NHE ear. The authors interpret this delay to be reflecting the lack of spectral information contained in degraded electrical signal produced by the CI which in turn leads to reduced access to semantic information and prolonged word evaluation (Finke et al. 2017).

Finke et al (2017) and Wedekind et al. (2021) both studied how the higher order processes in SSD CI users differ between the NHE and CI when stimulated independently. Analyzing each ear independently enables us to understand how the CI ear may be limited by the degraded electrical signal in comparison to the NHE. Currently, no ERP study in patients with SSD has compared whether the use of CI in conjunction with the NHE improves free field listening compared to the NHE alone.

### Current Study

In this study, we had two aims. Our primary aim was to evaluate the effect that the CI has in free field, comparing the difference between NHE alone and the integration of the CI and NHE. There are a few advantages to this approach, firstly, the presentation of words in free field is more representative of the real-world listening experience (compared to presentation of simpler sounds via direct connect or headphones). Secondly, measurement of neural and behavioural measures in the oddball task enables us to investigate how higher-order cortical processes are modulated by the additional electrical signal produced by the CI and whether this leads to a measurable change in task performance (e.g., reaction time [RT], RT variability and accuracy). Lastly, the contrast between FF-on and -off in free field allows us to characterise the interaction of signals between the NHE and CI. While it is well-documented that the addition of a CI facilitates hearing performance, previous research has identified the potential for interference between the electrical and organic signals in terms of neural processing (Finke et al. 2016), but this has yet to be shown to influence task behaviour.

As a secondary aim, we also sought to contrast neural responses and oddball task performance during monaural presentation to the CI and the NHE (via direct connect and headphones). This allows us to characterise the capabilities of CI alone and evaluate it against NHE performance, but also provides additional context to bridge ERP research using monaural and free field presentations. One challenge associated with translating between studies across different methods of presentation is that the experimental tasks can differ substantially across studies in a way that significantly impacts task performance. As such, it can be difficult to determine whether performance differences across studies are attributed to the method of presentation or task specific parameters (e.g., such as stimulus complexity, stimulus set, and background noise). By examining both monaural and free field presentations using the same task and participant sample, we can illustrate the degree of similarity between presentation methods on task performance. To further characterise the binaural hearing abilities of our SSD-CI participants, we also included clinical sound localisation and speech-in-noise comprehension tests.

## Method

### Participants

Ten SSD participants with a CI were recruited from the Fiona Stanley Hospital audiology department (Mean (SD)_age_ = 49(15.8) years, range 30 – 75 years, 7 female) (see Table 1). Participants were only asked to participate if they met all the inclusion criteria, no participant was excluded from the study. The main cause of deafness was idiopathic sudden sensorineural hearing loss in eight participants, with the other two participants losing their hearing due to Meniere’s disease and mumps. All participants had been using a MED-EL (Innsbruck, Austria) CI for at least one year with the majority of subject implanted with FLEX^28^ electrode array (two participants were implanted with a FLEX^26^ and FLEXSOFT electrode array). Participants recruited for this study used either a SONNET or RONDO speech processor. Mean unaided pure tone average (PTA) for the implanted ear was 90 dBHL (SD = 22.3). In the non-implanted ear, the mean PTA was 10 dBHL (SD = 3.9). Participants provided written informed consent prior to participating in the experiment. Ethics approval was obtained from the South Metropolitan Health Ethics Committee (reference number: 335).

**Table 1.** Demographic information of all participants. Age, Gender, Cochlear Implant (CI) Ear, Duration of deafness (years before implantation), Etiology (ISSNHL refers to idiopathic sudden sensorineural hearing loss), Cochlear implant (CI) ear pure tone audiometry (PTA) prior to implantation, Normal hearing ear (NHE) PTA, Type of implant, Electrode type, CI experience (time since implantation).

| Subject | Age | Gender | CI Ear | Duration of Deafness (years) | Etiology | CI PTA (db HL) | NHE PTA (db HL) | Type of Implant | Electrode | CI Experience (years) |
| --- | --- | --- | --- | --- | --- | --- | --- | --- | --- | --- |
| 1 | 75 | Male | Right | 0.8 | ISSNHL | 82 | 8 | Sonnet | FLEX28 | 5 |
| 2 | 46 | Male | Right | 2 | Meniere's Disease | 88 | 12 | Sonnet | FLEX28 | 2 |
| 3 | 38 | Female | Right | 4 | ISSNHL | 73 | 5 | Sonnet | FLEX28 | 3 |
| 4 | 30 | Female | Right | 25 | Mumps | 120 | 8 | Rondo | FLEX26 | 1 |
| 5 | 68 | Female | Right | 0.7 | ISSNHL | 63 | 16 | Sonnet | FLEX28 | 3 |
| 6 | 64 | Female | Left | 12 | ISSNHL | 83 | 9 | Sonnet | FLEX28 | 2 |
| 7 | 34 | Female | Right | 20 | ISSNHL | 56 | 6 | Rondo | FLEX28 | 2 |
| 8 | 56 | Female | Left | 0.9 | ISSNHL | 120 | 7 | Sonnet | FLEXSOFT | 8 |
| 9 | 34 | Female | Right | 7 | ISSNHL | 91 | 18 | Sonnet | FLEX28 | 3 |
| 10 | 55 | Male | Left | 30 | ISSNHL | 120 | 11 | Sonnet | FLEX28 | 5 |

### Speech Perception in Noise

The Bamford-Kowal-Bench Adaptive Speech-In-Noise test (Bench et al. 1979) was used to measure speech perception in noise. Three different spatial configurations were assessed in free field; 1) S0/N0: speech and noise presented from the front, 2) S_CI_/N_NHE_: speech presented to the CI and noise presented to the NHE and 3) S_0_/N_NHE_: speech presented from the front and noise to the normal hearing ear. Each configuration was assessed twice, once with the CI on and secondly without the CI. All speech perception in noise tests were conducted in an illuminated soundproof booth and block orders were counterbalanced across participants. All configurations were conducted by using a 3-speaker set up in free field placed 1 meter away at 0°. This set up has been used previously in our research group (Távora-Vieira et al. 2015, 2016; Wedekind et al. 2020; Wedekind et al. 2018; Wedekind et al. 2021).

### Localisation

Auditory Speech Sounds Evaluation Localisation Test was used to assess each SSD participant’s localisation of sound ability. Testing was conducted in a soundproof booth and a 4000 Hz narrowband sound was simultaneous presented through two speakers placed at −60 and 60 degrees from the participant. The two speakers presented the same 4000 Hz narrowband sound. To create a different interaural level difference, the presentation level through each speaker was randomised which created the illusion that the sound was coming from somewhere in between the two true speakers. All stimuli were presented at 60 dB HL at one loud speaker and depending on the interaural level difference (ILD), the other true speaker was presented at 60, 56, 40 or 30 dB. The software randomly picks an ILD from the folloiwing series: −30, −20, −10, −4, 0, +4, +10, +20, +30. This enabled 13 localisation points (two true speakers and 11 sham speakers) to be placed at 10-degree intervals in front of the subject (Tavora-Vieira et al. 2015). The two loudspeakers and 11 sham speakers were numbered from negative six to positive six and arranged into a semicircle with the participant sitting direclty in front of the loudspeaker numbered zero. The participant was required to report which one of the 13 speakers they heard the sounds from. Each response that the participant made was entered into a computer software which calculated the median values and root mean square (RMS) as a measure of a participant’s localisation ability. A lower RMS indicated better localisation ability. A total of 33 test items (narrowband noise) were randomly presented by a computer software. Stimuli intensity differences of −4, 0 and 4 dB were presented five times each and − 30, −20, −10, 10, 20, and 30 dB presented three times each. Auditory Speech Sounds Evaluation Localisation Test was used in the present study as it evaluates an individuals interaural level differences, which is thought to be the binaural tool that individuals use for sound localisation (Grantham et al. 2007). Each participant performed this test with and without the CI.

### Odd-Even Oddball Task

In this study, we used an oddball task where participants were presented with odd or even numbers and instructed to respond as soon as they heard an odd number (referred to as the ‘target’). Participant responded to target stimuli by pressing a button on a control pad. A set of eight stimuli were used (odd = one, three, five, nine; even = two, four, six, eight). Participants completed four blocks consisting of 180 trials, where target stimuli were presented on 20 % of trials (36 presentations) and standard stimuli were presented on 80% of trials (144 presentations). The duration of stimuli was ∼400 ms (average = 402 ms, range = 350 – 481 ms) with an onset-to-onset inter-stimulus interval of 1800 ms. Block order was counter-balanced across participants.

Speech files were attained from the National Acoustic Laboratories (Sydney, Australia) and were recorded by a mature female Australian speaker. These speech files were recorded with the purpose to be used in a telephone-based speech-in-noise test called ‘Telescreen’ (Dillon et al. 2016). Sounds were modified in audacity to reduce variation in duration between sounds while maintaining intelligibility.

In all four conditions, stimuli were presented at a calibrated intensity of ∼55 dB sound pressure level. The odd/even oddball paradigm was presented to the NHE via high-fidelity headphones (Audio-Technica ATH-M30x). For the direct connection to the CI, the stimuli were presented at a loud but comfortable level that was subjectively determined by the participant. During direct connect stimuli were directly sent to the CI via a cable from the computer’s 3.5mm audio jack, thereby bypassing the microphone of the CI. Free field conditions were presented using EDIFIER M1250 Multimedia Speakers that were positioned directly in front of the participant (Figure 1). See figure 1 for a schematic diagram of the four conditions. The experiment was programmed and delivered using Cogent 2000 and Psychtoolbox-3 functions in MATLAB. The behavioural and electrophysiological data were simultaneously collected from the oddball task.

**Figure 1.**
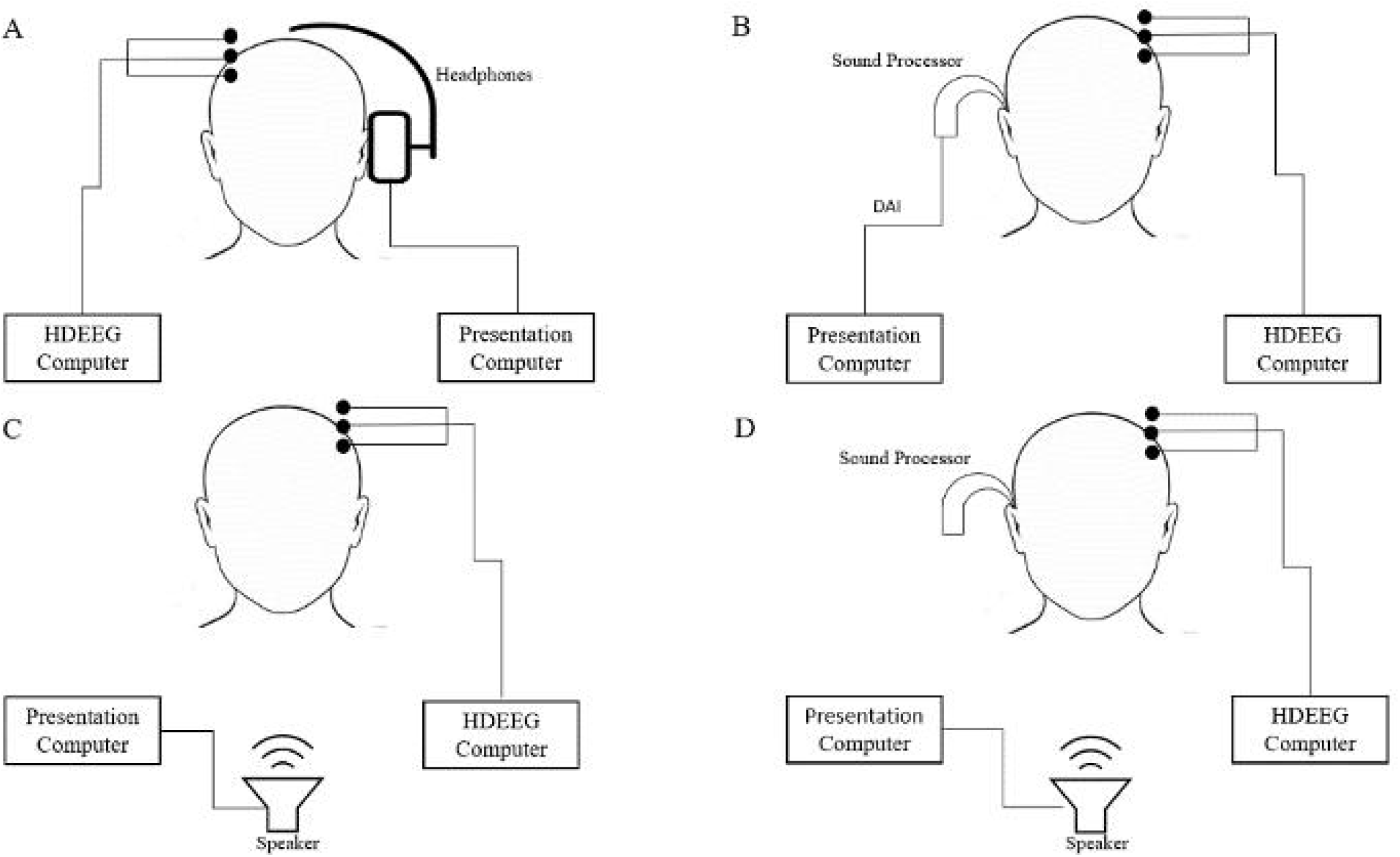
Schematic diagram depicting the four experimental conditions. **(A)** and **(B)** show the direct presentation of stimuli to the NHE and CI via headphones and direct connect, respectively. **(C)** and **(D)** show free field presentation with the FF-Off and FF-On, respectively.

### Acquisition and Pre-Processing of Electrophysiological Data

Electrophysiological data were recorded from 59+3 Ag/AgCl electrodes on a SpesMedica cap (10-5 layout) (SpesMedica ™ Genova, Italy) using the Micromed™ SD LTM EXPRESS system with Gilat Medical ERP software (Gilat Medical Research & Equipment Ltd, Karkur, Israel). Of the three additional electrodes, an electrode was placed on the infraorbital region of the right eye to record eye-movement, a reference electrode was placed on the middle of the chin, and a ground electrode was placed on the mastoid region contralateral to the CI. A sampling rate of 1024 Hz was used with a 40 Hz online low pass-filter. All electrode impedances were kept below 5 kΩ for the duration of the recording.

The data were pre-processed in MATLAB (ver. 2020a) using a semi-automated procedure consisting of functions from various plugins: EEGLAB (Delorme et al. 2004), PREP pipeline (Bigdely-Shamlo et al. 2015), AMICA (Palmer et al. 2011) and ICLabel (Pion-Tonachini et al. 2019). The data was filtered with a 1-30 Hz low pass filter was applied using the pop_eegfiltnew() function and were down-sampled to 250 Hz. The pop_clean_rawdata() function was used to identify noisy channels. The pop_interp() function was used to interpolate bad channels using the spherical method. The clean_asr() function was used to correct for artefacts using the artefact subspace reconstruction method. Epochs segments were extracted for each trial, spanning −200 to 1000 ms from stimulus-onset. Independent component analysis of the data was conducted using AMICA (2000 iterations) (Palmer et al. 2011). ICLabel was used to identify independent components associated with artifacts with > 70% confidence (i.e., eye movement, muscle, heart, line noise, and channel noise). The data were baseline corrected at the trial-level by subtracting the mean amplitude within pre-stimulus interval (−200 to 0 ms) from the entire ERP waveform. Trials with activity exceeding 100 µV were excluded from further analysis. Incorrect trials, defined as: standard trials with a response, target trials without a response, or target trials with RTs < 50 or > 1500 ms were omitted from all further calculation and analyses. After screening, the mean (SD) number of target trials retained per participant were NHE: 33.3(3.30) 92.5%, CI: 23.1(6.10) 64.2%, FF-Off: 31.2(8.07) 86.7% and FF-On: 32.5(4.55) 90.3%. For standard trials, mean retained trials were NHE: 136.8(18.59) 95%, CI: 116.4(40.35) 80.8%, FF-Off: 125.8(39.11) 87.36%, FF-On: 131.6(23.41) 91.4%.

### Measurement of Event-Related Potentials

We measured N1 and P2 mean amplitudes at the trial-level over 40 ms time-windows centred around the respective peaks of the N1 and P2 at the frontocentral electrode (FCz) where peak amplitudes were maximal. Measurement intervals were determined by identifying negative/positive peak of the grand-average waveform (across standard and target trials) between 100-250 ms and 200-350 ms respectively, and measuring ±20 ms around the peaks. Separate measurement windows were calculated for each presentation to account for temporal jitter. The measurement windows for N1 were NHE: 136-176 ms, CI: 148-188 ms, FF-Off: 144-184 ms, FF-On: 144-184 ms. The measurement windows for P2 were NHE: 244-284 ms, CI: 228-268 ms, FF-Off: 252-292 ms, FF-On: 252-292 ms.

We measured N2N4 and P3b mean amplitude at the trial-level over 40 ms time-windows at left fronto-central (FC3) and parietal midline (Pz) respectively, where the target-standard N2N4 and P3b effects were maximal. Measurement windows were determined by identifying negative/positive peaks of the grand-averaged ‘target-minus-standard’ difference waveforms between 250-450 ms and 400-650 ms respectively, and measuring ±20 ms around the peaks. Separate measurement windows were calculated for each presentation to account for temporal jitter and morphological variations. The measurement windows for N2N4 were NHE: 436-476 ms, CI: 436-476 ms, FF-Off: 428-468 ms, FF-On: 436-476 ms. Measurement windows for P3b were NHE: 556-596 ms, CI: 520-560 ms, FF-Off: 536-576 ms, FF-On: 636-676 ms).

### Behavioural Data

We examined task performance by measuring RT, RT variability and target accuracy. Based on the distribution of RTs, we accepted RTs within 50 to 1500 ms on target trials only. Target accuracy was calculated as the number of target trials with a response within the accepted window. RT variability was calculated as the standard deviation of RTs within each condition.

### Statistical Analysis

Statistical analyses were conducted using R statistics and R Studio software (R 2013). Direct (NHE vs. CI) and free-field presentations (FF-On vs FF-Off) were analysed separately. Linear mixed models analysis was conducted using the ‘lme’ function from the ‘nlme’ package (Pinheiro et al. 2022). For behavioural measures (Localisation, Speech In Noise Comprehension Test, RT, RT variability and Accuracy), we included Ear (CI/NHE or FF-on/FF-off) as a fixed-effect, and intercepts for participants were modelled as a random effect. For electrophysiological measures (N1, P2, N2N4 and P3b), we also included Trial-type (Standard, Target) as an additional fixed-effect, interacting with the effect of Ear.

We presented the results as F-values using the ‘anova’ function. Follow-up pairwise comparisons were conducted using ‘emmeans’ and ‘contrast’ functions from the ‘emmeans’ package (Lenth et al. 2020). We presented the results of these pairwise categorical comparisons as t-ratios (mean difference estimate divided by standard error) and p-values for multiple comparisons were corrected using the ‘Holm’ method. Values and error bars for plots presented in the results were also derived from the ‘emmeans’, which produces estimated marignal means – a summary projected from the linear mixed model which considers the varaibiltiy across trial-observations and the random-effects, and is more representative of the statistical analysis than raw means. In our description of ERP amplitudes, we use the term ‘enhance’ to describe an increase in amplitude in the direction respective of the polarity for each ERP component.

## Results

### Localisation and Speech Perception in Noise Test

On the Localisation test (Fig. 2A), linear mixed model analyses revealed a significant effect of CI on RMS error (*F*(1,8) = 36.36, *p* = 0.003*), indicating that sound localisation performance was superior with FF-On compared to FF-Off (*M*(*SD*) = 21.11(8.27) vs. 47.78(13.59) degrees error, *est. mean diff*. (*SE*) = 26.7(4.42)).

**Figure 2.**
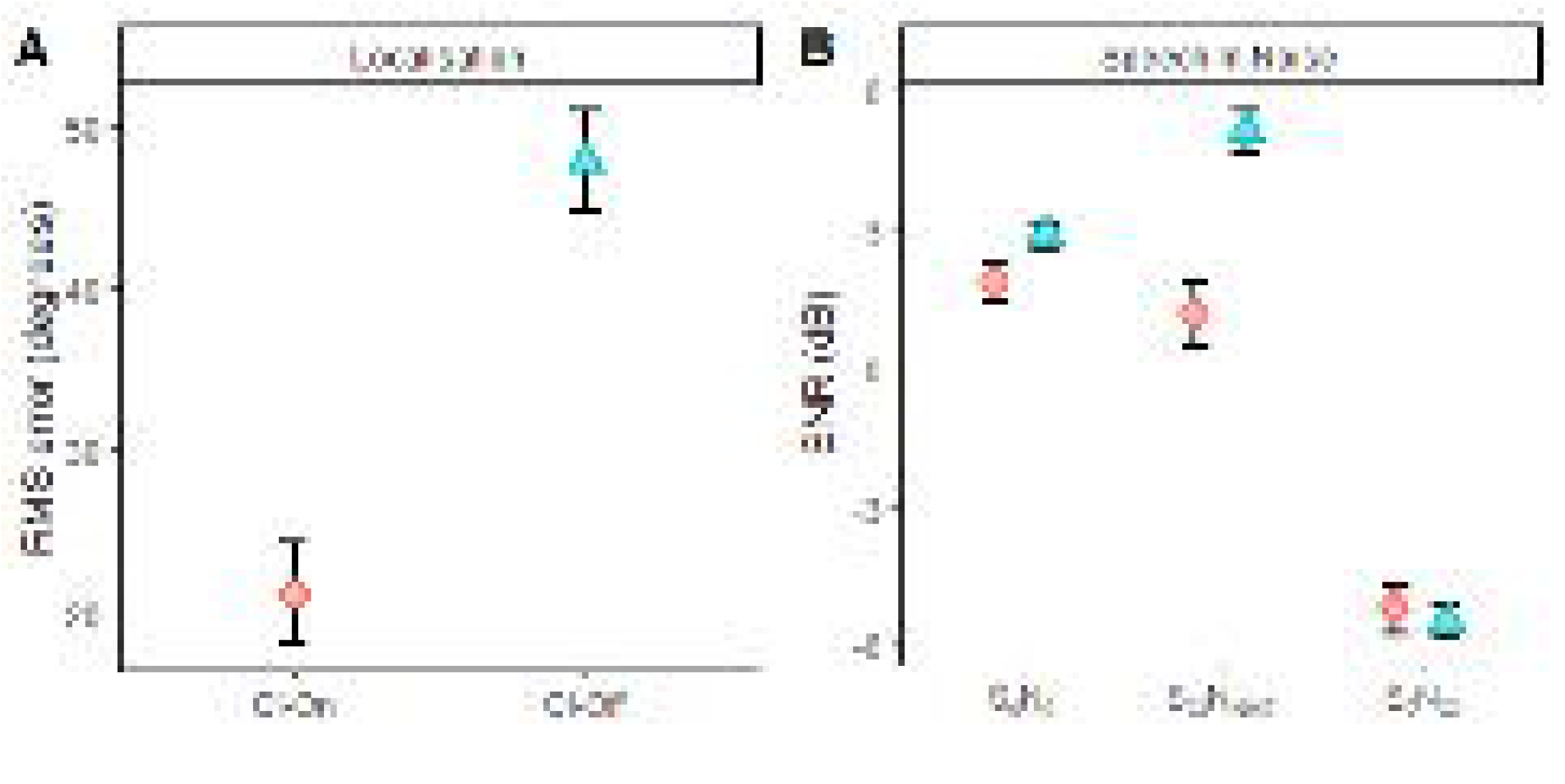
Estimated marginal mean values (with error bars depicting the estimated standard error of the mean) for **(A)** Localisation and **(B)** Speech Perception in Noise tests.

On the Speech Perception in Noise test (Fig. 2B), linear mixed model analysis revealed a significant main effects of test-condition (*F*(2,45) = 189.57, *p* < .001*) and CI (*F*(1,25) = 15.56, *p* = .003*), as well as a significant test-condition × CI interaction (*F*(2,45) = 10.24, *p* = .002*). Follow-up pairwise comparisons revealed that the signal-to-noise ratio (SNR) was significantly lower with CI-On during S_CI_N_NHE_ (M(SD) = 1.15(3.07) vs. 5.10(2.31) dB, *t-ratio*(45) = 5.80, *p* < .001*, *est. diff*. (*SE*) = 3.95(0.68) dB), but SNRs were comparable during S_0_N_0_ (*M*(*SD*) = 1.15(3.07) vs. 2.85(1.13) dB, *t-ratio*(45) = 1.47, *p* = .297, *est. diff*. (*SE*) = 1(0.68) dB) and S_0_N_CI_ (*M*(*SD*) = −5.20(2.16) vs. −5.50(1.79) dB, *t-ratio*(45) = 0.44, *p* = 0.662, *est. diff*. (*SE*) = 0.3(0.68) dB).

### Reaction Time

For reaction time (Fig. 3A), we observed significantly longer RTs to target stimuli when presented to the CI via direct connect compared to the NHE presentation via headphones (*F*(1,580) = 84.71, *p* < .001*, *M*(*SD*) = 842.35(128.35) vs. 720.67(94.18) ms, *est. mean diff*. (*SE*) = 116(12.6) ms). In free field, linear mixed analyses indicated a statistically significant increase in RTs during FF-On compared to FF-Off (*F*(1,676) = 5.38, *p* = .021*, *M*(*SD*) = 737.91(94.03) vs. 709.04(103.51), *est. mean diff*. (*SE*) = 27.4(11.8) ms).

**Figure 3.**
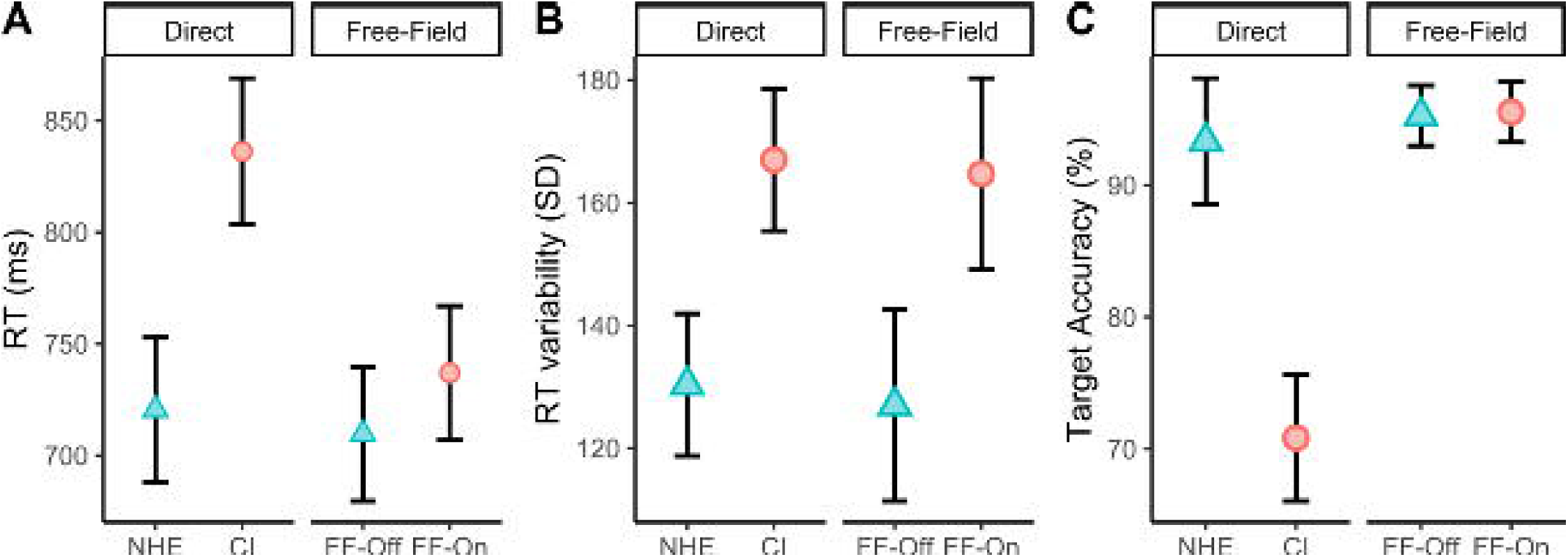
Estimated marginal mean values (with error bars depicting the estimated standard error of the mean) for **(A)** Reaction Time and **(B)** Reaction Time Variability and **(C)** Target Accuracy in the Oddball Task.

### Reaction Time Variability

For RT variability (Fig. 3B), we observed that the standard deviation of RTs was greater during direct CI compared to direct NHE presentation (*F*(1,9) = 9.75, *p* = .012*, *M*(*SD*) = 167.01(31.35) vs. 130.34(41.31), *est. mean diff*. (*SE*) = 36.7(11.7) ms). Interestingly, in free field, RT variability was also greater during FF-On compared to FF-Off (*F*(1,9) = 10.46, *p* = 010*, *M*(*SD*) = 164.71(50.81) vs. 127.00(47.90), *est. mean diff*. (*SE*) = 37.7(11.7) ms).

### Accuracy

For accuracy (Fig. 3C), our results followed the same pattern as RT – we observed a lower accuracy on target trials during direct CI compared to direct NHE presentation (*F*(1,9) = 13.15, *p* = .006, *M*(*SD*) = 70.83(19.21) vs. 93.33(9.46) %, *est. mean diff*. (*SE*) = 22.5(6.2) %). However, in free field, target accuracy was very high and did not differ between FF-On and FF-Off conditions (*F*(1,9) = 0.06, *p* = .811, *M*(*SD*) = 95.56(6.17) vs. 95.27(8.18)%, *est. mean diff*. (*SE*) = 0.28(1.13) %).

### N1 mean amplitude

During direct presentation, a significant main effect of Ear was observed (*F*(1,3083) = 4.83, *p* = .028*), indicating that overall N1 amplitude was more negative on NHE compared to CI (*M*(*SD*) = −1.04(0.88) vs. −0.89(0.88) µV, *est. mean diff*. (*SE*) = 0.16(0.11) µV). There was no significant main effect of Trial-type (*F*(1,3083) = 1.40, *p* = 0.237) or Ear × Trial-type Interaction (*F*(1,3083) = 0.13, *p* = .718).

During free-field presentation, a significant main effect of Ear was observed (*F*(1,3198) = 7.96, *p* = .005*), indicating that overall N1 amplitude was more negative on FF-on compared to FF-off (*M*(*SD*) = −1.34(0.80) vs. −1.20(0.97) µV, *est. mean diff*. (*SE*) = 0.22(0.12) µV). There was no significant effect of Trial-type (*F*(1,3198) = 0.67, *p* = .412) or Ear × Trial-type interaction (*F*(1,3198) = 0.67, *p* = .412).

### P2 mean amplitude

During direct presentation, a significant main effect of Ear was observed (*F*(1,3083) = 67.31, *p* < .0001*), indicating that P2 amplitude was overall more positive for NHE compared to CI (*M*(*SD*) = 1.09(1.44) vs. 0.48(0.71) µV, *est. mean diff*. (*SE*) = 0.83(0.12) µV). There was also a significant effect of Trial-Type (*F*(1,3083) = 4.08, *p* = .043*) indicating that P2 amplitude was more positive on standard compared to target trials, but no significant Ear × Trial-Type interaction was observed (*F*(1,3083) = 1.45, *p* = .229) (Figure 4A & 4B).

**Figure 4.**
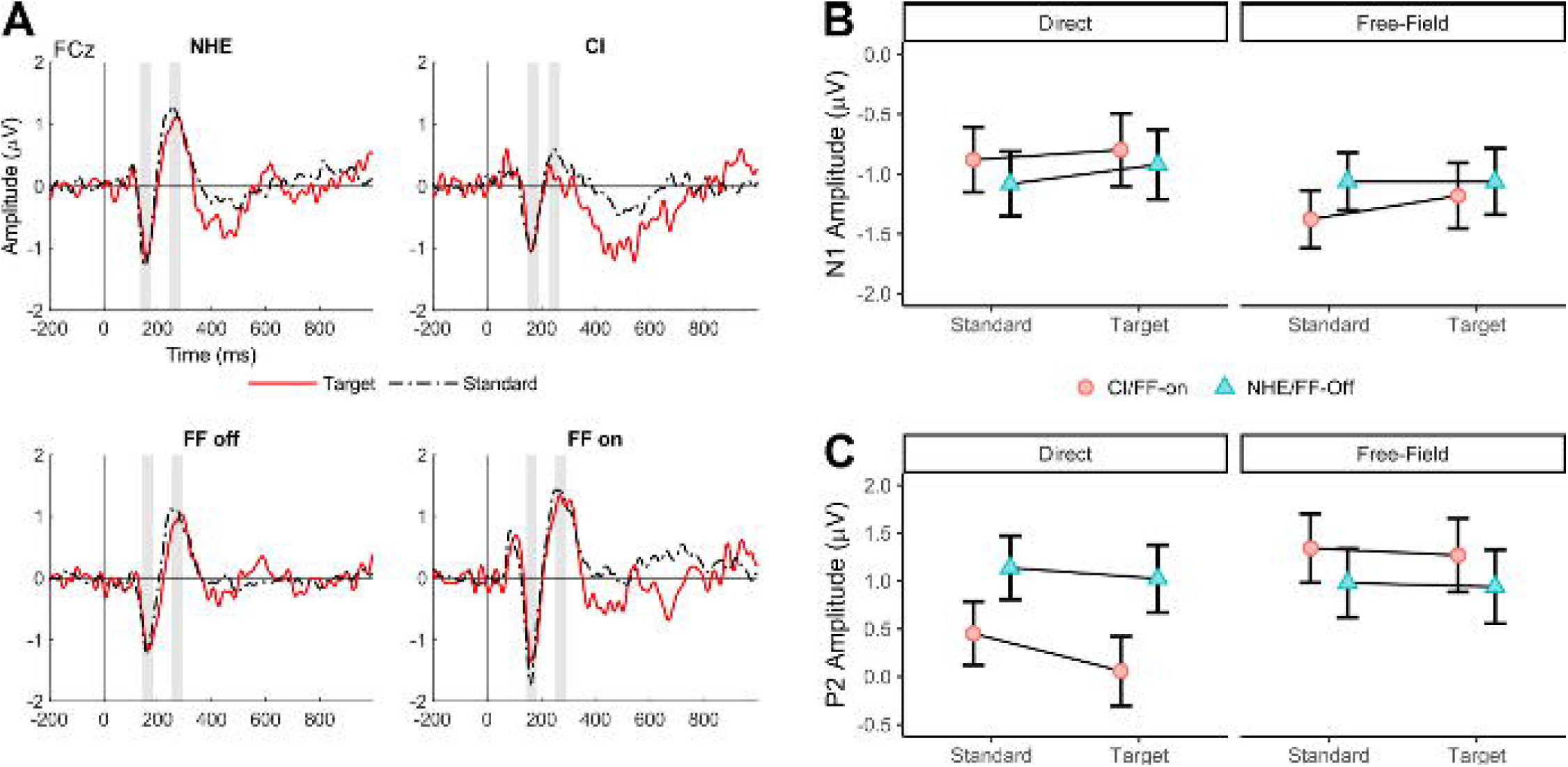
(**A**) Grand mean ERP waveforms for each stimulus (Standard, Target) and presentation configuration (NHE, CI, Free-Field-Off, Free-Field-On) with shaded areas depicting N1 and P2 measurement intervals. (**B**) Grand mean values (with within-subject standard error bars) for N1 and (**C**) P2 mean amplitude.

During free-field presentation, a significant effect of Ear was observed (*F*(1,3198) = 12.37, *p* = .0004*). However, the main effect of Trial-Type (*F*(1,3198) = 0.20, *p* = .652) and Ear × Trial-type interaction *F*(1,3198) = 0.18, *p* = 0.893) were not statistically significant. These results indicate that P2 amplitude was more positive for FF-on compared to FF-off (*M*(*SD*) = 1.34(1.36) vs. 0.92 (1.03) µV, *est. mean diff*. (*SE*) = 0.34(0.13) µV), but P2 amplitude did not differ significantly between standard and target trial (Figure 4C).

### N2N4 Mean Amplitude

During direct presentation, the linear mixed model revealed significant effects of Ear (*F*(1,3083) = 6.96, *p* = .0.008*) and Trial-Type (*F*(1,3083) = 77.53, *p* < .0001*), but the Ear × Trial-type interaction was not statistically significant (*F*(1,3083) = 1.64, *p* = .200). These results indicate that N2N4 amplitude was significantly enhanced on target compared to standard trials (*M*(*SD*) = −1.31(1.47) vs. −0.23(0.24) µV, *est. mean diff*. (*SE*) = 0.95(0.11) µV), and that there was an overall negative offset for NHE compared to CI (M(SD) = −0.46(0.50) vs. −0.28(0.51) µV, *est. mean diff*. (*SE*) = −0.29(0.11) µV), but the size of the N2N4 effect (target-standard difference) was not significantly different for NHE and CI. During free-field presentation, there was a significant main effect of Trial-type (*F*(1,3198) = 73.92, *p* < .0001*) and Ear × Trial-type interaction *F*(1,3198) = 8.28, *p* = .004*). However, the main effect of Ear was not statistically significant *F*(1,3198) = 0.60, *p* = .439). The results indicate that N2N4 amplitude was significantly enhanced on target compared to standard trials (*M*(*SD*) = −0.95(1.34) vs. 0.07(0.33) µV, *est. mean diff*. (*SE*) = 0.94(0.11) µV), and follow-up comparisons showed that size of the N2N4 effect was larger for FF-on (*t-ratio*(3198) = 8.16, *p* < .0001*, *est. mean diff*. (*SE*) = 1.25(0.15) µV) compared to FF-off (*t-ratio*(3198) = 3.96, *p* < .0001*, *est. mean diff*. (*SE*) = 0.62(0.16) µV) (Figure 5A & 5B).

**Figure 5.**
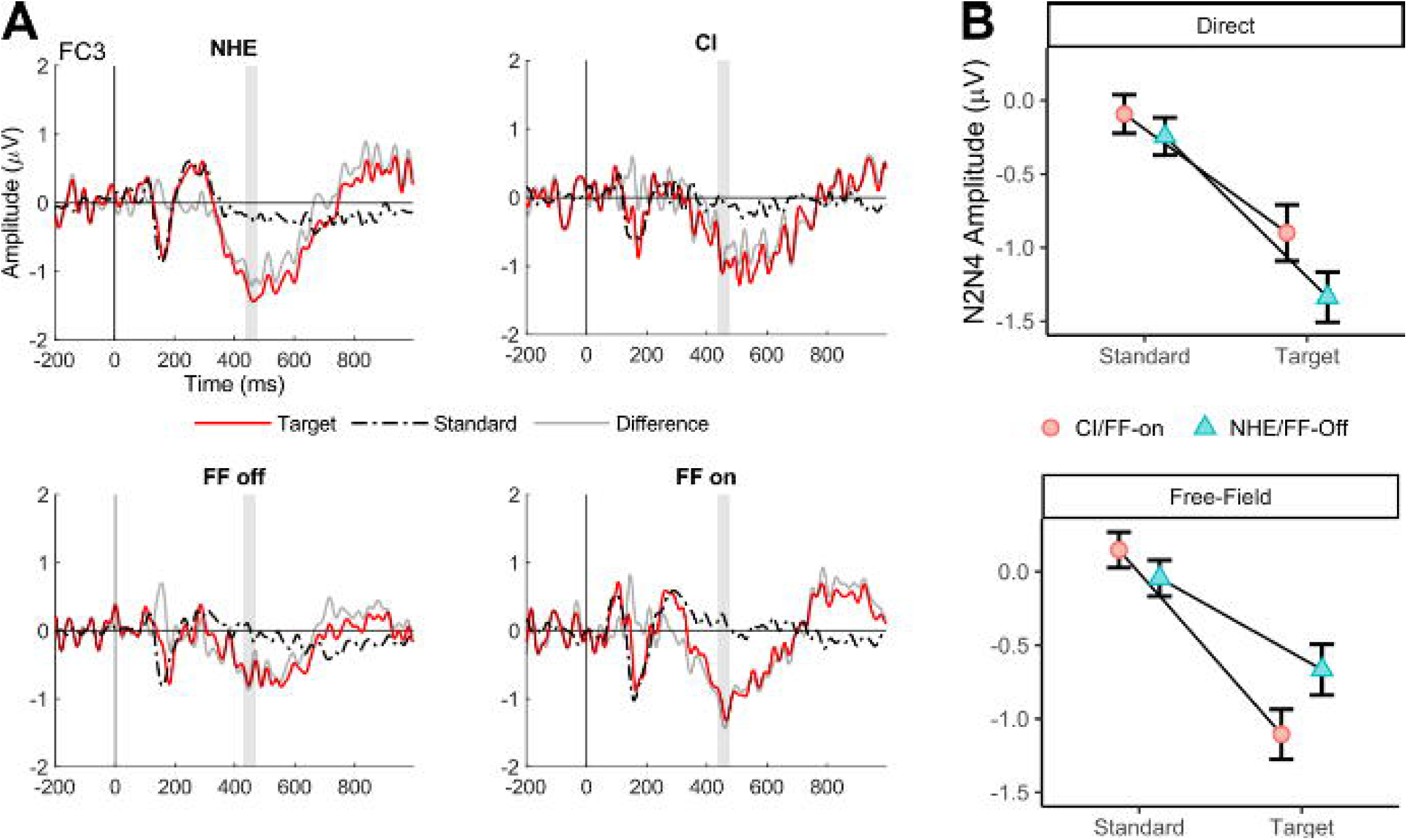
**(A)** Grand average waveforms for all conditions condition using a single left frontocentral electrode (FC3). The grey shaded area indicates the time window used to measure the N2N4. **(B)** Mean amplitude of N2N4 for all four conditions for standard and target trials. Error bars depict within subject standard error bars.

### P3b Mean Amplitude

During direct presentation, a significant main effect of Trial-type was observed (*F*(1,3083) = 27.17, *p* < .0001*). However, no significant main effect of Ear (*F*(1,3083) = 0.78, *p* = .376) and the Ear × Trial-type interaction (*F*(1,3083) = 0.58, *p* = .447) were observed. These results indicate that P3b was more positive on target compared to standard trials (*M*(*SD*) = 1.31(0.89) vs. 0.55(0.67) µV, *est. mean diff*. (*SE*) = 0.74(0.15) µV), but there was no significant difference in overall amplitude or the size of the P3b effect between Ears.

During free-field presentation, we observed a significant main effect of Trial-type (*F*(1,3198) = 28.06, *p* < .0001*) and a significant Ear × Trial-Type interaction (*F*(1,3198) = 5.37, *p* = .021*). However, the main effect of Ear was not statistically significant (*F*(1,3198) = 0.57, *p* = .449). The results indicate that P3b was more positive on target compared to standard trials (*M*(*SD*) = 0.89(0.43) vs. 0.16(0.65) µV, *est. mean diff*. (*SE*) = 0.64(0.12) µV), and follow-up analyses indicated that size of the P3b effect was larger on FF-on (*t-ratio*(3198) = 5.41, *p* < .001*, *est. mean diff*. (*SE*) = 0.93(0.17) µV) compared to FF-off (t-ratio(3198) = 2.05, *p* = .040*, *est. mean diff*. (*SE*) = 0.36(0.18) µV) (Figure 6A & 6B).

**Figure 6.**
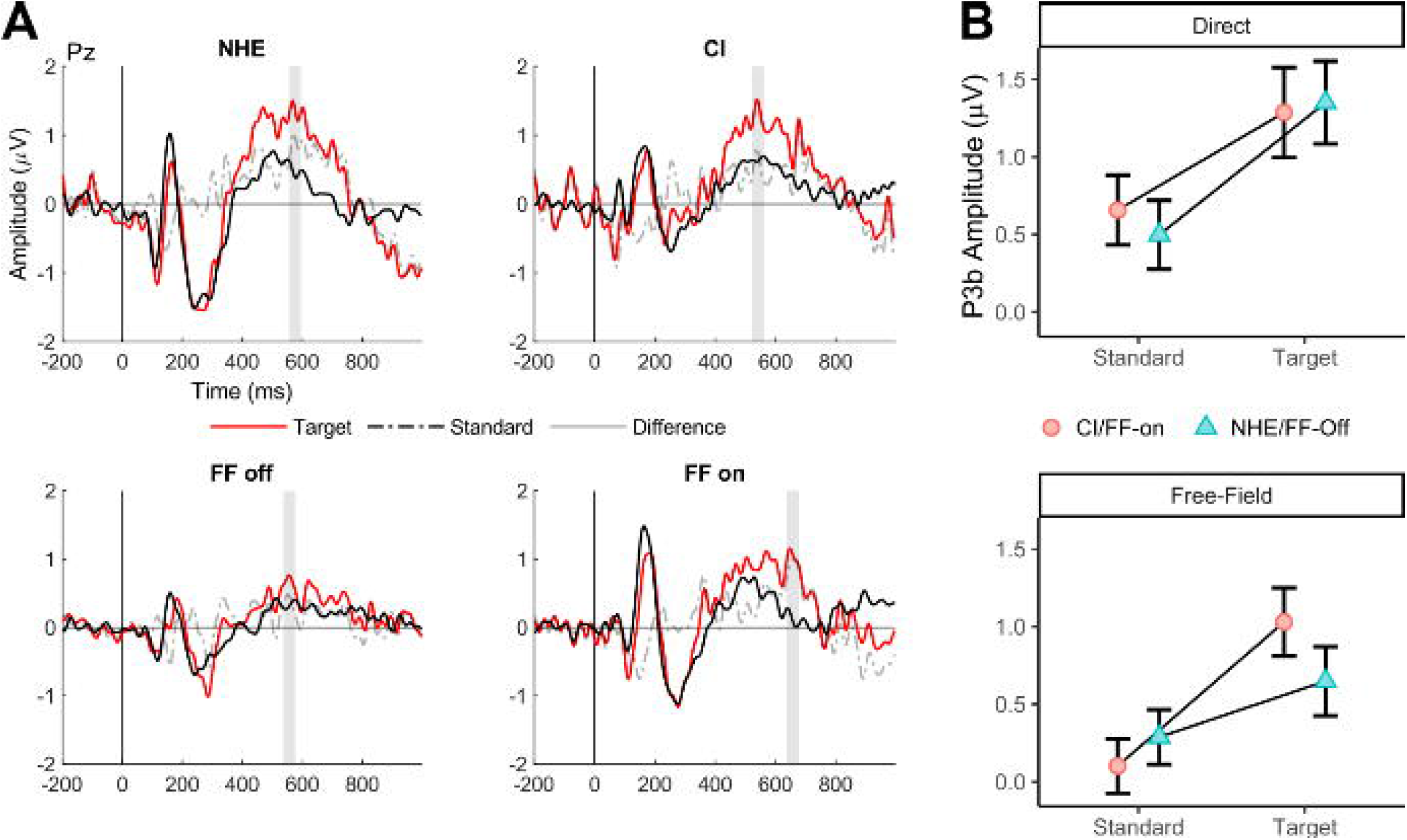
**(A)** Grand average waveforms for all four conditions using a single parietal electrode (Pz). The grey shaded area indicates the time window used to measure the P3b. **(B)** Mean amplitude of P3b for all conditions for standard and target trials.

## Discussion

In the current study, we sought to systematically examine how hearing in various monaural and bilateral configurations (direct and free field) impacts oddball task performance and neural responses associated with higher-order cognitive processing. We also characterised hearing ability in our SSD CI sample using sound localisation and speech in noise comprehension tests and identified that performance in both tests improve significantly when the CI was switched on. In the semantic oddball task, we found that participants were capable of discriminating target and standard stimuli with the CI alone. However, CI performance was sub-optimal compared to the NHE. In free-field, target accuracy was similarly high in both FF-on and FF-off. Interestingly, RTs were more variable, and slightly but significantly delayed during FF-on compared to FF-off. Overall neural responses showed identifiable N1 and P2 peaks, with standard-target differences in N2N4 and P3b in each condition. Collectively, these results demonstrate that the semantic oddball task is sensitive to the presence of CI under both direct and free-field configurations, highlight both the capabilities and limitations of the CI alone compared to the NHE, and important insights about optimising task design when assessing bilateral hearing in SSD-CI individuals.

### Task performance: CI is less capable, and participants rely on NHE during bilateral hearing

In monoaural hearing conditions, we observed delayed RTs and lower target accuracy when the stimuli were delivered directly to the CI. This finding is consistent with previous monaural studies and likely reflects the differences in the quality of electrical signal coming from the CI, compared to the NHE. The size of the RT difference was ∼100 ms, which falls in between previous observations. When comparing the NHE to the CI, evaluating each ear independently, Wedekind et al. (2021) observed differences of ∼40 ms during tone-discrimination and Finke et al. (2016) observed differences of ∼150 ms during word-discrimination. This difference is unlikely attributed to the delay from CI, which was identified to be 1.5 ms (Zirn et al. 2015) and are likely due to delays in decision time increases with stimulus and/or task complexity.

With respect to accuracy, we observed ∼70 % correct on target trials for direct CI, whereas Finke et al. (2016) reported slightly lower hit rates (∼65%). Comparing our task with Finke et al. (2016), we used mono-syllable words with a small word set (8 words), whereas they used a di-syllable words with a much larger word set (35 words) – therefore RT and accuracy effects in our study could be due to either or a combination of these differences.

During free field presentation, we observed high accuracy for both FF-on and FF-off (∼95%), similar to the NHE (∼92%) suggesting that participants were largely relying on the NHE to perform the task. Interestingly, we also identified that RTs were more variable but significantly delayed (∼27 ms) for FF-On when compared to FF-Off. This finding is consistent with previous speculation that asymmetries in signals produced by the CI and NHE may lead to interference (Finke et al. 2016; Wedekind et al. 2020), and the presence of this interference confirms that CI signals are being processed by the brain. Although the absence of overall improvement is counter intuitive given that performance on sound localisation and speech-in-noise tests improved significantly with CI, these results are better understood considering the task’s simplified design. In a simple task with no binaural cues, the addition of CI did not provide any extra useful information, limiting the possible headroom for improvement and explaining why performance does not exceed the NHE alone. Increasing task difficulty by using a larger word-set and more complex sounds in future studies may increase the likelihood of detecting such an effect.

### Electrophysiological Results: Early ERPs enhanced during NHE

For direct conditions, early neural responses reflecting stimulus detection (N1 and P2) were larger in amplitude for NHE, suggesting that sound signals from the NHE were more salient. This highlights that the degraded electrical signal produced by the CI fails to encapsulate all the spectral information that the NHE provides. This is consistent with previous reported that N1-P2 amplitude were smaller in the SSD CI users when compared to normal hearing controls (Legris et al. 2020). During free-field, early neural responses were larger in amplitude during FF-On compared to FF-Off, and likely reflects the effect of additional signal input from the CI facilitating lower-level detection reflected by the N1, and enhancing higher attentional-orientation processes reflected by the P2 (Wedekind et al. 2018). In Figure 6 (Pz), an early positivity occurring at the same latency to the frontal N1 also shows a similar pattern of results across conditions. One possibility is that this positivity is an inversion of the N1, reflecting the opposing side of same dipole.

For later ERPs, we observed enhanced N2N4 and P3b mean amplitudes to target compared to standard stimuli across all hearing configurations, highlighting that higher-order discriminatory and evaluative processes reflected by these ERPs were able to differentiate between stimuli, consistent with previous oddball studies (Delle-Vigne et al. 2015; Kotchoubey et al. 2001; Li et al. 2019; Tomé et al. 2015). We observed larger differences in N2N4 and P3b between standard and target trials during FF-on compared to FF-off. However, we did not observe any significant differences in the size of the N2N4 and P3b effects between CI and the NHE conditions.

While the absence of N2N4 effects between direct conditions contrast with Finke et al. (2016) who reported enhanced N2N4 amplitudes during the direct connect condition, the observation of a larger N2N4 effect during FF-on is in line with their results. In Finke et al. (2016), the authors attributed the increase in N2N4 during CI to lexical processing, reflecting greater effort to retrieve and match sounds with the mental lexicon. It is possible that elevated N2N4 during FF-on may reflect increased effort needed to resolve lower-level uncertainties associated with the addition of the CI signal, making speech more difficult to understand (Cope et al. 2017; Obleser et al. 2007)

However, the observed increase in P3b during FF-on compared to FF-off is more difficult to interpret. It is possible that this enhancement may also reflect increased effort to resolve lower-level uncertainties, similar to the N2N4. However, this interpretation conflicts with Finke et al. (2016) who reported reduced P3b during CI may reflect difficulty evaluating and classifying living and non-living words. While our ERP results are largely consistent with previous studies, discrepancies in the direction of P3b effects and the relationship between neural responses and task performance across direct and free-field configurations (i.e., large behavioural difference for direct but no N2N4/P3b differences during direct, and the opposite for free-field) highlight that these brain-behaviour relationships are complex and are likely indirect in nature. Highlighting the complexities of interpreting ERPs, early ERP differences may have downstream effects which alter the interpretating of later ERPs. For example, in the scenario where there are early N1 and/or P2 ERP differences between conditions, the absence of subsequent N2N4 amplitude differences between direct NHE and CI conditions may not unequivocally indicate that the underlying discriminatory processes are indifferent. It is possible that these processes have been modulated by earlier differences but are not captured by amplitude measures. However, these effects are not yet understood as they are difficult to measure.

### Conclusion, Limitations and Future Directions

In this study, we sought to examine how bilateral hearing in SSD-CI users compares to hearing in monaural configurations. Previous work has established that listening with the CI alone was less efficient compared to the NHE, but research has yet to systematically explore the interactions that take place during bilateral hearing. In our data, we found that oddball performance was similar between bilateral hearing (FF-on and FF-off) compared to the NHE alone, demonstrating a reliance on the NHE, which was more than capable for performing the required task.

In a simple task with no binaural cues, the addition of the CI did not provide any additional information that the NHE could provide – explain why performance does not exceed the NHE alone. Although the addition of CI did not benefit performance during FF-on, CI signals were still being processed by the brain – reflected by increases in RT variability during CI and FF-on conditions as well as enhanced P2 amplitudes and N2N4 and P3b effects during FF-off. It is clear that CI and NHE signals are being used together, but current task (1) did not provide binaural cues (such as spatial audio) that would allow to exceed the NHE and (2) use of a limited set of mono-syllable words may have produced ceiling effects in RT/accuracy preventing any benefit in binaural hearing to be observed.

Considering these factors, future studies examining binaural hearing in SSD CI users should employ a design that (1) uses spatial cues and (2) is more difficult, either by using a larger word set or by presenting background noise. With respect to the neural response, we failed to report ERP differences between NHE and CI conditions, despite clear behavioural differences. While this may be explained by using a simple task leading to ceiling effects in the ERP response, it remains to be seen whether these differences can be observed in more complex and difficult tasks.

Other avenues for future research in the CI ERP literature is to directly contrast neural responses during unilateral and bilateral presentations. Although we examined ERPs in both configurations, the use of different presentation modes (e.g., we used combination of direct connect, headphones and speakers) does limit the comparability of these signals. One way of reducing these differences is to simulate free field listening using direct connect, which has been previously demonstrated (Arnoff et al., 2010). This avenue would be most suited for bilateral CI samples.

